# Entomological surveys and insecticide susceptibility profile of *Aedes aegypti* during the dengue outbreak in Sao Tome and Principe in 2022

**DOI:** 10.1101/2024.01.07.574582

**Authors:** Basile Kamgang, João Acântara, Armel Tedjou, Christophe Keumeni, Aurelie Yougang, Anne Ancia, Françoise Bigirimana, Sian Clarke, Vilfrido Santana Gil, Charles Wondji

## Abstract

**Background:** The first dengue outbreak was reported in Sao Tome and Principe in 2022. This study was undertaken aiming to establish the typology of *Aedes* larval habitats, the distribution of *Ae. aegypti* and *Ae. albopictus,* the related entomological risk and the susceptibility profile of *Ae. aegypti* to insecticides for a better evidence-informed response.

**Methodology/Principal Findings:** Entomological surveys were performed in all the seven health districts of Sao tome and Principe during the dry and rainy seasons in 2022. WHO tube and synergist assays using piperonyl butoxide (PBO) and diethyl maleate (DEM) were carried out and the genotyping of F1534C/V1016I/V410L mutations in *Ae. aegypti*. *Aedes aegypti* and *Ae. albopictus* were found in all seven health districts of the country with high prevalence of *Ae. aegypti* in the most urbanised district, Agua Grande. Both *Aedes* species bred mainly in used tyres, discarded tanks and water storage containers. In both survey periods, the Breteau (BI > 50), house (HI > 35%) and container (CI > 20%) indices were higher than threshold established by WHO to indicate high potential risk of dengue transmission. *Aedes aegypti* sample was susceptible to all insecticides tested except dichlorodiphenyltrichloroethane (DDT) (9.2% mortality), bendiocarb (61.4% mortality) and alpha-cypermethrin (97% mortality). A full recovery was reported in *Ae. aegypti* resistant to bendiocarb after pre-exposure to synergist PBO. Only one *Ae. aegypti* specimen was found carrying F1534C mutation.

**Conclusions/Significance:** These findings revealed at high potential risk for dengue transmission throughout the year, with the bulk of larval breeding occurring in used tyres, water storage and discarded containers. Most of the insecticides tested remain effective to control *Aedes* vectors in Sao Tome except DDT and bendiocarb. These data underline the importance to raise community awareness and to implement routine dengue vector control strategies to prevent further outbreaks in Sao Tome and Principe, and elsewhere in the subregion.

**Author Summary:** During the first dengue outbreak reported in Sao Tome and Principe in 2022, entomological investigations were undertaken aiming to establish the typology of *Aedes* larval habitats, the distribution of *Ae. aegypti* and *Ae. albopictus,* the related entomological risk and the susceptibility profile of *Ae. aegypti* to insecticides for a better evidence-informed response. The results revealed the presence of *Ae. aegypti* and *Ae. albopictus* in all seven health districts of the country with high prevalence of *Ae. aegypti* in the most urbanised district, Agua Grande. Both *Aedes* species bred mainly in used tyres, discarded tanks and water storage containers suggesting a good waste management and improving water supply system could help to reduce *Aedes* densities and the risk of dengue transmission. Analyses also revealed that most of the insecticides tested remain effective to control *Aedes* vectors in Sao Tome except dichlorodiphenyltrichloroethane and bendiocarb. These findings revealed at high potential risk for dengue transmission throughout the year and underline the importance to raise community awareness and to implement routine dengue vector control strategies to prevent further outbreaks in Sao Tome and Principe, and elsewhere in the subregion.

## Background

Dengue is the most important mosquito-borne viral disease in the world in terms of mortality and morbidity. Indeed, one modelling estimate indicates 390 million dengue virus infections per year, of which 96 million (67–136 million) manifest clinically [1]. Dengue is caused by the dengue virus (DENV) belonging to the *Flavivirus* genus and *Flaviviridae* family. This virus is transmitted to vertebrates including humans by the bite of infected female *Aedes* species mosquitoes. Formerly, dengue was considered scarce in Africa probably due to the similarity of symptoms with other infectious diseases like malaria. In countries with endemic malaria, it was therefore often misdiagnosed. However, during the past two decades, increasing number of cases of dengue have been reported across the continent with major outbreaks in some countries including Gabon [2], Burkina Faso [3], and Angola [4]. In addition, a study realised in Douala Cameroon between July and December 2020 demonstrated that ∼13% of acute febrile patients presenting in health facilities in Douala Cameroon are due to dengue [5] suggesting that the dengue prevalence in Africa can be higher than expected. The first case of dengue was reported in Sao Tome and Principe in April 2022, reaching a cumulative total of 1,200 cases by March 2023, with the peak of the outbreak in June 2022. Since there is no specific treatment nor efficient vaccine against dengue, the control of dengue outbreaks therefore remains reliant mainly on vector control through the destruction of *Aedes* larval habitats and the application of insecticides to either treat larval habitats or control adults [6]. The implementation of this control strategy requires a good knowledge of the vectors involved in the outbreak.

Both epidemic prone vectors of dengue, *Aedes aegypti* and *Ae. albopictus* are found in Central Africa but these species have different origins. *Aedes aegypti* originated from African forests [7] whereas *Ae. albopictus* originates from South-East Asia forest [8], but has since spread globally and invaded all the continents during the last four decades. This invasive species was first reported in Central Africa in early 2000s in Cameroon [9] and has rapidly spread in almost all central African countries including Sao Tome and Principe where it was first reported in 2016 [10]. Studies performed in Central Africa on larval ecology of both species showed that they are colonising the same types of larval habitats composed mainly of used tyres, discarded tanks and water storage containers [11]. However, in the cities where both species are found, *Ae. aegypti* prefers larval habitats located in the downtown neighbourhoods with high building density while *Ae. albopictus* is mostly found in peri-urban or rural areas surrounded by vegetation [12–14]. The emergence of insecticide resistance in *Aedes* mosquitoes can seriously compromise vector control using insecticides as demonstrated in some countries [15, 16]. Data generated in numerous Central African countries reveals significant variation of insecticide resistance according to *Aedes* species tested, the origin of mosquito and the type of insecticide [17–22]. Both major mechanisms, metabolic resistance and target site resistance, generally found involved in insecticide resistance in *Aedes* mosquitoes, were suspected as the main causes of resistance in Central Africa. Indeed, knock down resistance (*kdr*) mutations, F1534C, V410L and V1016G were detected in *Ae. aegypti* with high frequency for 1534C [19]. Some cytochrome P450 genes were found over expressed in *Ae. aegypti* and *Ae. albopictus* from Central Africa [19].

In Sao Tome and Principe data on *Aedes* in this regard are very scarce. To fill this knowledge gap, this entomological investigation aimed to assess the typology of larval habitats, the entomological risk using Stegomyian indices, the current distribution of *Ae. aegypti* and *Ae. albopictus* as well as the resistance profile of *Ae. aegypti* to insecticides.

## Methods

### Study area

Surveys were carried out in Sao Tome and Principe, an island country of Central Africa. It consists of two islands, Sao Tome, the bigger one subdivided into six health districts: Agua Grande, Lobata, Me-zochi, Caue, Cantagalo and Lemba, and Principe, the smaller one (RAP: Região Autonóma do Príncipe) representing one health district (Figure 1). The population of both islands is estimated to 200,000 inhabitants. The climate is humid tropical characterised by two seasons: a long rainy season of nine months duration from September to May and a shorter dry season from June to August; mean annual rainfall is 1,382 mm. Mean temperatures vary a few degrees throughout the year, ranging between 22°C and 26°C. The lowest average temperatures occur during the dry season, while the rainy season experiences higher temperatures (https://climateknowledgeportal.worldbank.org/country/sao-tome-and-principe/climate-data-historical).

**Figure 1.**
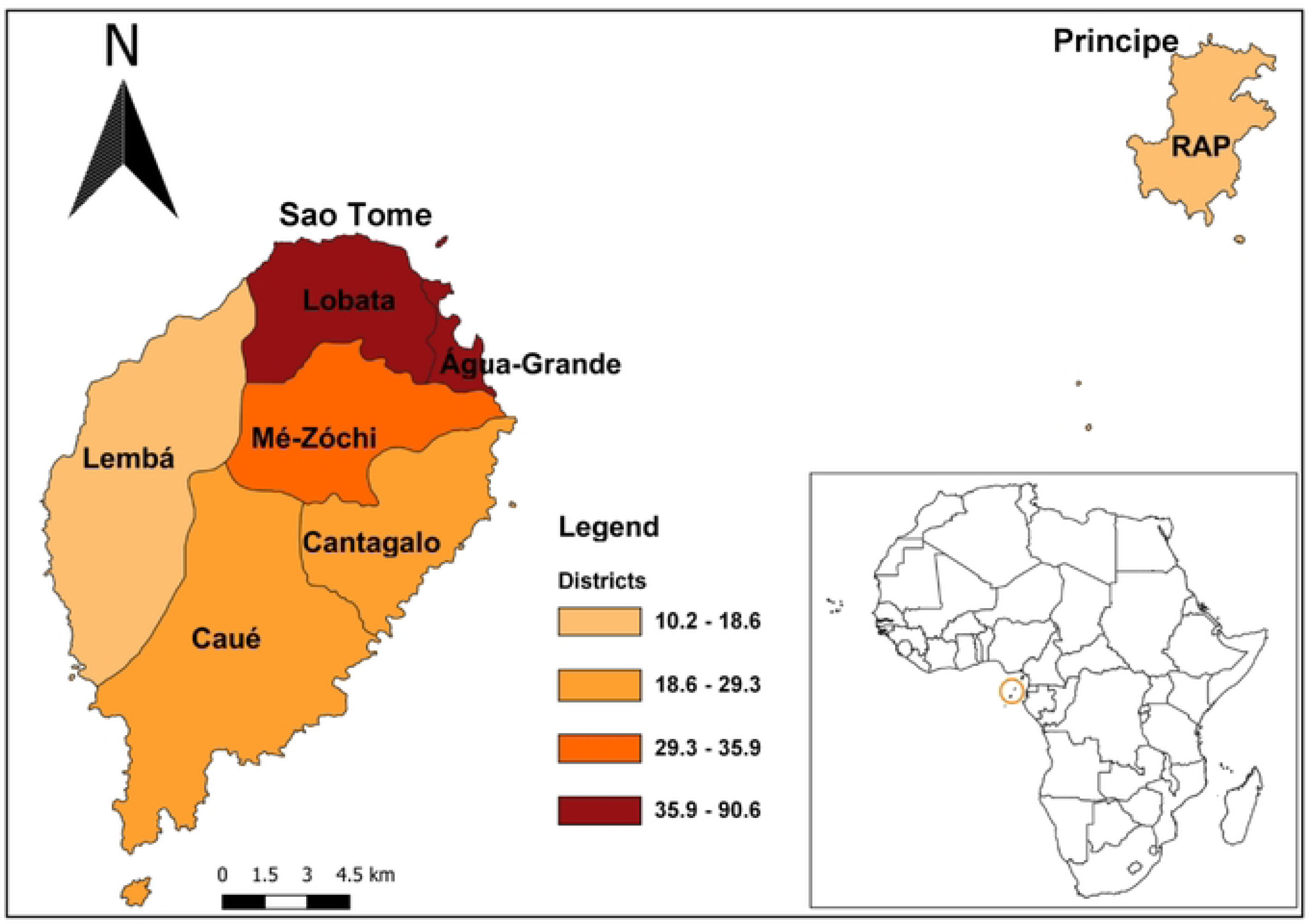
Map of Sao Tome and Principe indicating the seven health districts. Colors indicate population density by health district.

### Sampling and entomological surveys

Entomological investigations were carried out in June and November 2022, corresponding to the dry and rainy seasons respectively. Nationwide surveys were undertaken in clusters of randomly selected dwellings in each of seven health districts of the country. In each district, a minimum of three clusters were randomly selected; each cluster consisted of 15 dwellings per neighbourhood. During the surveys, each selected dwelling and its surroundings was inspected to record all natural and/or artificial containers with water (potential larval habitat), and those containing at least one larvae or pupae (positive larval habitat). On basis of the nature, the source and use of the water, potential larval habitats were classified into three categories: domestic, peri-domestic, and natural. Domestic containers (e.g. storage tanks) were defined as human-filled receptacles, while peri-domestic (e.g. discarded tanks, used tyres), and natural receptacles (e.g. rock and tree holes, leaf axils) were those filled by rain [13]. Larvae and pupae found per container were collected and transported to the insectary and isolated from predators such as *Lutzia tigripes* and reared to adults. Emerged adults were morphologically identified alive using a suitable taxonomic key [23, 24]. Adult mosquitoes identified as *Ae. aegypti* or *Ae. albopictus* were pooled and reared until obtaining the adults of G1 generation used to performed adult bioassays to insecticide. The number of immature stages of each species was estimated from the proportion of emerging adults of each species. During this field investigation all the discarded tanks identified were destroyed and water storage containers were treated with larvicide (Fig 2). Local entomologists from different health districts in Sao Tome and Principe were trained on the identification of *Aedes* larval habitats and its destruction or treatment. Advice was also provided to the population on how to avoid and eliminate *Aedes* larval habitats in their environment.

**Figure 2.**
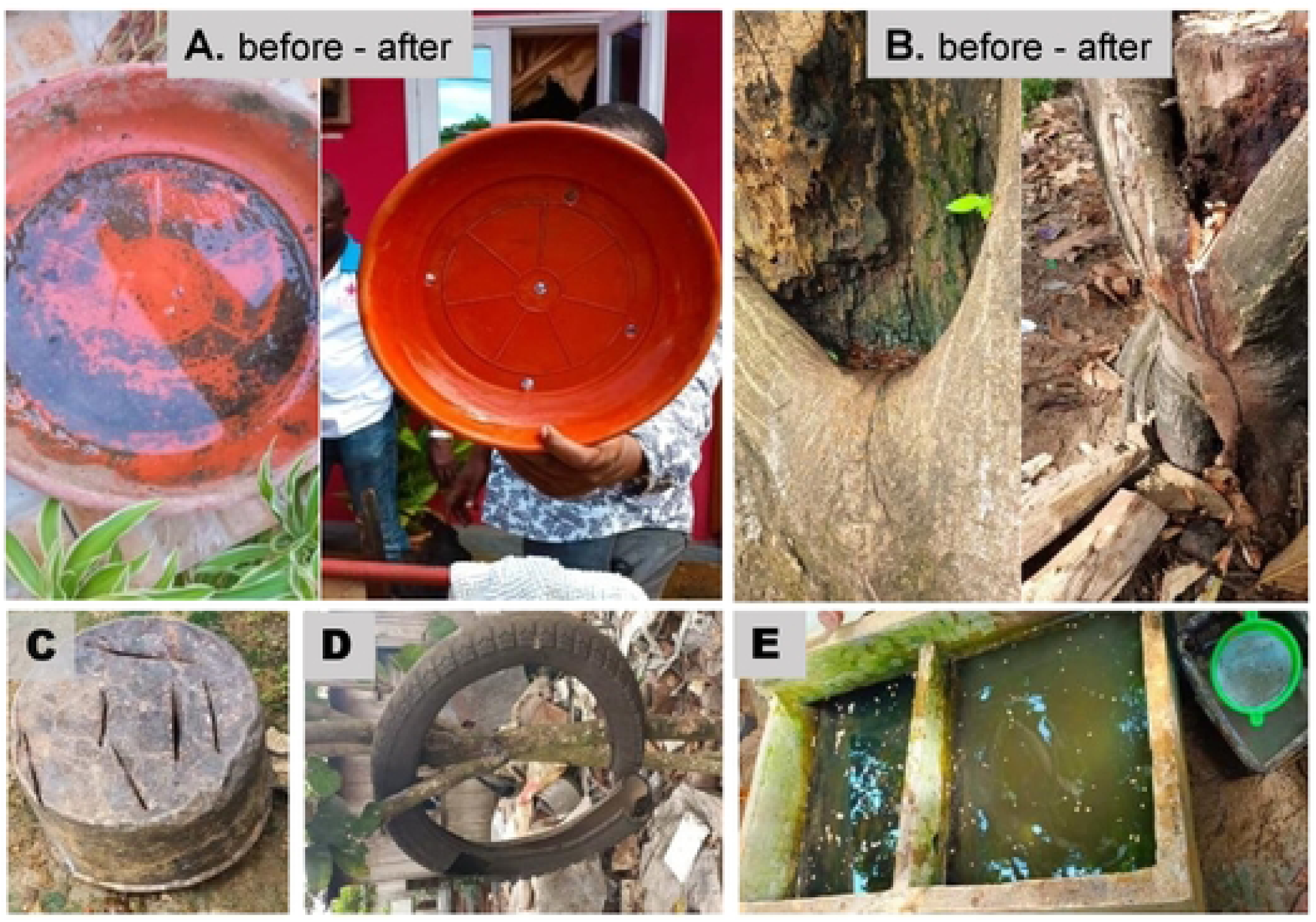
Pictures indicating some examples of action taken on the field during the dengue outbreak to control *Aedes* larvae. A, Destroying plastic saucer of flower pot by perforation with holes; B, Destroying of tree hole (natural larval habitat) by creating drainage channel; C, Destroying of discarded tanks by perforation; D, Destroying used tyres by cutting; E, Water storage container (cement tanks) treated with *Bacillus thuringiensis israelensis* (*Bti*).

### Entomological indices

The level of infestation was estimated using traditional Stegomyia indices including Breteau index (BI, the number of positive containers per 100 prospected houses), house index (HI, the percentage of houses infested), and container index (CI, percentage of positive containers). Estimated reference thresholds of HI, BI, and CI established by WHO for dengue and yellow fever transmission risk were applied: whenever HI > 35%, BI > 50, and CI > 20%, the location is considered as high risk of urban transmission of yellow fever virus, whereas HI < 4%, BI < 5 and CI < 3%, is considered to indicate a low risk of disease transmission [25]. Similarly, the categories of low HI < 0.1%, medium HI 0.1%-5% and high HI > 5% were established for classifying risk of dengue transmission [26].

### Insecticide resistance evaluation in *Ae. aegypti*

For this study *Ae. aegypti* Benin strain was used as the reference full susceptible laboratory strain. Bioassays were performed according to WHO protocol using 2-5 days old G1 generation. Four replicates of 20-25 females per tube were exposed to 0.03% deltamethrin, 0.40% permethrin, 0.25% pirimiphos-methyl, 0.05% alphacypermethrin, 0.1% bendiocarb, 1% fenitrothion and 4% dichlorodiphenyltrichloroethane (DDT) for 1 hour. Mortality was recorded 24 hours later and mosquitoes alive or dead after exposure were stored in RNA later or silica gel, respectively. The resistance status was defined as follows: susceptible (mortality rate between 98–100%), probable resistance (mortality rate between 90–97%), and resistant (mortality rate < 90%) [27].

### Adult synergist assay with PBO and DEM

To evaluate the potential role of oxidases and glutathione S-transferases (*GSTs*) in metabolic resistance mechanisms, synergist assays with 4% piperonyl butoxide (PBO) and 8% diethyl maleate (DEM) were performed. Two-five-days-old adults were pre-exposed for one hour to PBO or DEM impregnated papers and after that immediately exposed to the selected insecticide. Mortality was scored 24 hrs later and compared to the results obtained with each insecticide without synergist according to WHO standards [27]. The comparison of mortality rates after pre-exposure of mosquitoes to synergist and without pre-exposure to synergist was performed using Chi-square test.

### Knockdown resistance (*kdr*) genotyping in *Ae. aegypti*

Thirty specimens of *Ae. aegypti* from G0 were genotyped for three different *kdr* mutations: V1016I, V410L and F1534C, chosen because these mutations have been described as involved in pyrethroid resistance of *Ae. aegypti* mosquito [28, 29]. These mutations have also been previously reported in Central Africa [19]. Based on Moyes et al., review, F1534C and V410L are associated to insecticide resistance and V1016I is associated to insecticide resistance when combined to other *kdr* mutations [30]. Genomic DNA was extracted using the Livak protocol [31], and genotyping of the V1016I, V410L and F1534C mutations was performed by real-time melting curve quantitative PCR [32]. Each PCR reaction was performed in a 21.5 μL volume PCR tube containing 2 μL of DNA sample, 10 μL of SYBR® Green (SuperMix), and 1.25 μL of each primer. The amplification conditions were set as follow: 95°C for 3 min, followed by 40 cycles of (95°C for 20 s, 60°C for 1 min and 72°C for 30 s) and then final steps of 72°C for 5 min, 95°C for 1 min, 55°C for 30 s and 95°C for 30 s.

## Data analysis

Variables defined as categorical variables were summarised by percentages and confidence interval, and numeric variables by means and standard deviations; and compared using the Chi squared and Kruskal Wallis tests, respectively. All statistical analyses were performed with R version 4.2.1 and RStudio version 2023.03.0 (R Core Team, 2018), and *p*-value < 0.05 was considered statistically significant.

## Results

### Pre-imaginal infestation

In total we investigated 173 and 241 houses in 22 neighbourhood clusters across six health districts of Sao Tome, during the dry and rainy seasons respectively. In addition, 42 houses in four neighbourhoods were surveyed in Principe (RAP) during the dry season only.

In Sao Tome, out of 253 potential larval habitats for *Aedes* inspected during the dry season 123 (50.2%) were found positive for *Ae. aegypti* and/or *Ae. albopictus,* while 385 of 624 potential containers (61.7%) were positive in the rainy season (Table 1). The Stegomyia indices estimated in dry season were significantly lower than those calculated in rainy season: house index (HI 41.5 vs 69.3), Breteau index (BI 74.3 vs 163.5), and container index (CI 50.2 vs 63.2) respectively (Fig 3A). However, in both seasons, all the Stegomyia indices calculated were high and above the thresholds established by WHO (Fig 3A) to indicate potential high risk for dengue transmission. When analyses were performed according to district, HI varied significantly during the dry season (ꭓ^2^= 28.117, df = 5, *p* < 0.00001) with highest index (67.7) in Mezochi and lowest index (9.5) in Cantagalo (Fig 3C). A similar pattern was observed for the Breteau Index with the highest index (122.5) in Agua Grande and lowest (22.7) in Caue (ꭓ^2^ = 26.064, df = 5, *p* <0.00001). However, no significant difference was found for the CI. When similar analyses were performed for data collected during the rainy season, there was no significant variation in the three indexes between the districts (Fig 3D).

**Figure 3.**
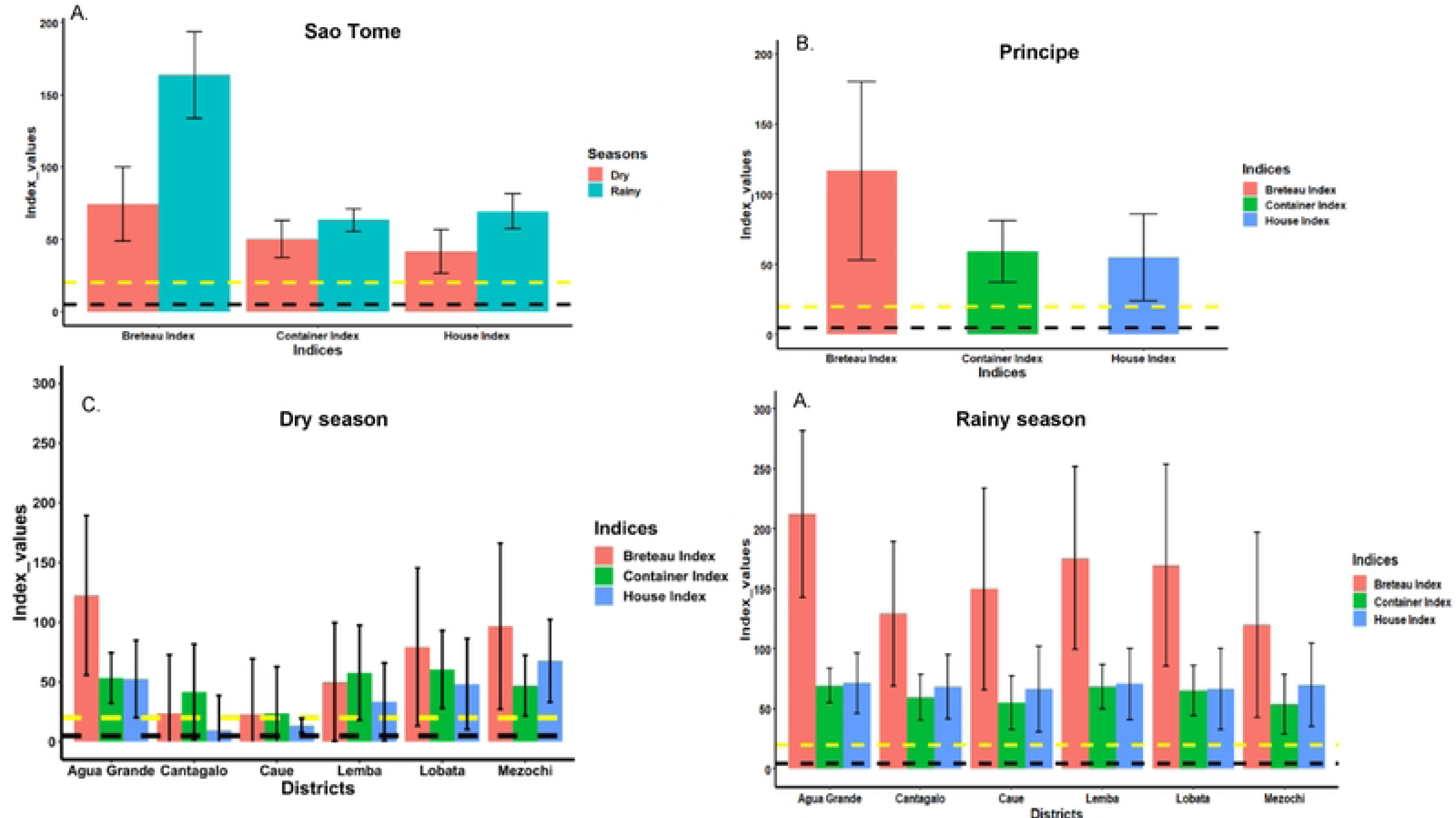
Stegomyia indices calculated in Sao Tome and Principe. A, Indices calculated in Sao Tome according to season. B, Indices calculated in Principe during the dry season. C, Indices calculated during the dry season in Sao Tome per health district; D, Indices calculated during the rainy season in Sao tome per health district.

**Table 1:** Container type infested with larvae and or pupae of Aedes spp. in rainy and dry seasons in Sao Tome, 2022.

| Container | Dry season (June 2022) |  |  |  | Rainy season (November 2022) |  |  |  |
| --- | --- | --- | --- | --- | --- | --- | --- | --- |
|  | N inspected | N positive | % positive | Relative contribution: % positive containers | N inspected | N positive | % positive | Relative contribution: % of positive containers |
| **Domestic** | **74** | **35** | **47.3** | **27.5** | **164** | **79** | **48.2** | **20.5** |
| Water storage | 48 | 22 | 45.8 | 17.3 | 130 | 61 | 46.9 | 15.8 |
| Flower pot | 13 | 9 | 69.2 | 7.1 | 23 | 11 | 47.8 | 2.9 |
| Animal drinking pot | 13 | 4 | 30.8 | 3.1 | 11 | 7 | 63.6 | 1.8 |
| **Peridomestic** | **161** | **86** | **53.4** | **67.7** | **430** | **283** | **65.8** | **73.5** |
| Used tyres | 67 | 35 | 52.2 | 27.6 | 163 | 111 | 68.1 | 28.8 |
| Discarded tanks | 70 | 34 | 48.6 | 26.8 | 150 | 96 | 64 | 24.9 |
| Plastic cover | 0 | 0 | - | 0 | 29 | 20 | 69 | 5.2 |
| Tin cans | 0 | 0 | - | 0 | 23 | 12 | 52.2 | 3.1 |
| Miscellaneous | 24 | 17 | 70.8 | 13.4 | 65 | 44 | 67.7 | 11.4 |
| **Natural** | **4** | **1** | **25** | **0.8** | **30** | **23** | **76.7** | **6.0** |
| Bamboo | 0 | 0 | - | 0 | 11 | 11 | 100 | 2.8 |
| Coconut shell | 0 | 0 | - | 0 | 7 | 4 | 57.1 | 1.0 |
| Snail shell | 0 | 0 | - | 0 | 7 | 3 | 42.9 | 0.9 |
| Tree holes | 1 | 1 | 100 | 0.8 | 2 | 2 | 100 | 0.5 |
| Rock holes | 0 | 0 | - | 0 | 3 | 3 | 100 | 0.8 |
| Axil of plant | 3 | 0 | 0 | 0 | 0 | 0 | - | 0 |
| **Overall** | **253** | **127** | **50.2** | **100 (n=127)** | **624** | **385** | **61.7** | **100 (n=385)** |

N inspected, Number of containers with water inspected; N positive, Number of containers found with immature stages of *Aedes* spp.; % positive, Percentage of each type of container found with immature stages of *Aedes* spp.; Relative contribution: Percentage of all positive containers found with immature stages of *Aedes* spp.

In Principe, all three indexes calculated during the dry season (HI= 54.8, BI = 116.7 and CI = 59.0) were high and superior to the threshold established by WHO for potential high transmission risk of dengue (Fig 3B).

### Typology and prevalence of larval habitat

During the entomological investigations in Sao Tome and Principe, all three categories of *Aedes* larval habitats were found: human-filled domestic (water storage container, flower pot and animal drinking pot), rain-filled peri-domestic (used tyres, tin cans, car wrecks and miscellaneous) and natural (tree holes, rock holes, snail shells, coconut shell, axil of plant). The number of each type of potential container and prevalence of containers infested with immature stage of *Aedes* for each island is presented in Tables 1 (Sao Tome) and Table 2 (Principe).

**Table 2:** Container type infested with larvae and/or pupae of *Aedes* spp. in dry season (June 2022) in Principe.

| Container | N inspected | N positive | % positive | Relative contribution: % positive containers |
| --- | --- | --- | --- | --- |
| **Domestic** | **65** | **37** | **56.9** | **67.3** |
| Water storage | 61 | 34 | 55.7 | 61.8 |
| Flower pot | 2 | 1 | 50.0 | 1.8 |
| Animal drinking pot | 2 | 2 | 100 | 3.7 |
| **Peridomestic** | **25** | **18** | **72.0** | **32.7** |
| Used tyres | 12 | 9 | 75.0 | 16.4 |
| Discarded tanks | 10 | 8 | 80.0 | 14.5 |
| Miscellaneous | 3 | 1 | 33.3 | 1.8 |
| **Natural** | **0** | **-** | **-** | **0.0** |
| **Overall** | **90** | **55** | **61.1** | **100 (n=55)** |

N inspected, Number of containers with water inspected; N positive, Number of containers found with immature stages of *Aedes* spp.; % positive, Percentage of each type of containers found with immature stages of *Aedes* spp.; Relative contribution: Percentage of all positive containers found with immature stages of *Aedes* spp.

In Sao Tome the most numerous potential larval habitats for *Aedes* spp. inspected were used tyres followed by discarded tanks and water storage containers, independent of the season. These containers were also found to be the most infested with larvae and /or pupae of *Aedes* spp. (Table 1). High infestation rates were observed in all domestic and peridomestic containers inspected on Sao Tome, however the greater number of discarded containers in the peridomestic environment results in these contributing most to *Aedes* vector breeding in the locations surveyed, in both seasons. An increased contribution of natural larval habitats was also noted during the rainy season, with the prevalence of natural containers infested with *Aedes* spp. of 5.98 % compared to just 0.79% in dry season. In Principe where inspections were only carried out during the dry season, domestic water storage containers were the most numerous potential larval habitats and also the main receptacles found infested (61.82%) with immature stages of *Ae. aegypti* and/or *Ae. albopictus*. This was followed by used tyres and discarded tanks (Table 2).

### Distribution of *Aedes aegypti* and *Ae. albopictus* in Sao Tome and Principe

In Sao Tome, 1,224 and 2,995 specimens of *Aedes* spp. were morphologically identified during the dry and rainy seasons respectively. These specimens were composed of 55.47% *Ae. aegypti* versus 44.53% *Ae. albopictus* in the dry season, and 53.46% *Ae. aegypti* versus 46.54% *Ae. albopictus* in the rainy season. Thus overall, *Ae. aegypti* was the most prevalent *Aedes* species in both seasons in Sao Tome. However, when analysed according to the district, *Ae. albopictus* was found most prevalent in Caue and Cantagalo, while *Ae. aegypti* was the predominant species in Agua Grande, irrespective to the season (Figure 4). Nevertheless, in Lemba, Lobata and Mezochi the relative prevalence of the two *Aedes* species varied according to the season, with a significantly higher prevalence of *Ae. albopictus* seen in Lemba and Mezochi during dry season (Figure 4). In Principe, where investigations were carried out only during the dry season, a total of 126 individuals of *Aedes* spp. were identified comprising 48 % *Ae. aegypti* and 52% *Ae. albopictus*.

**Figure 4.**
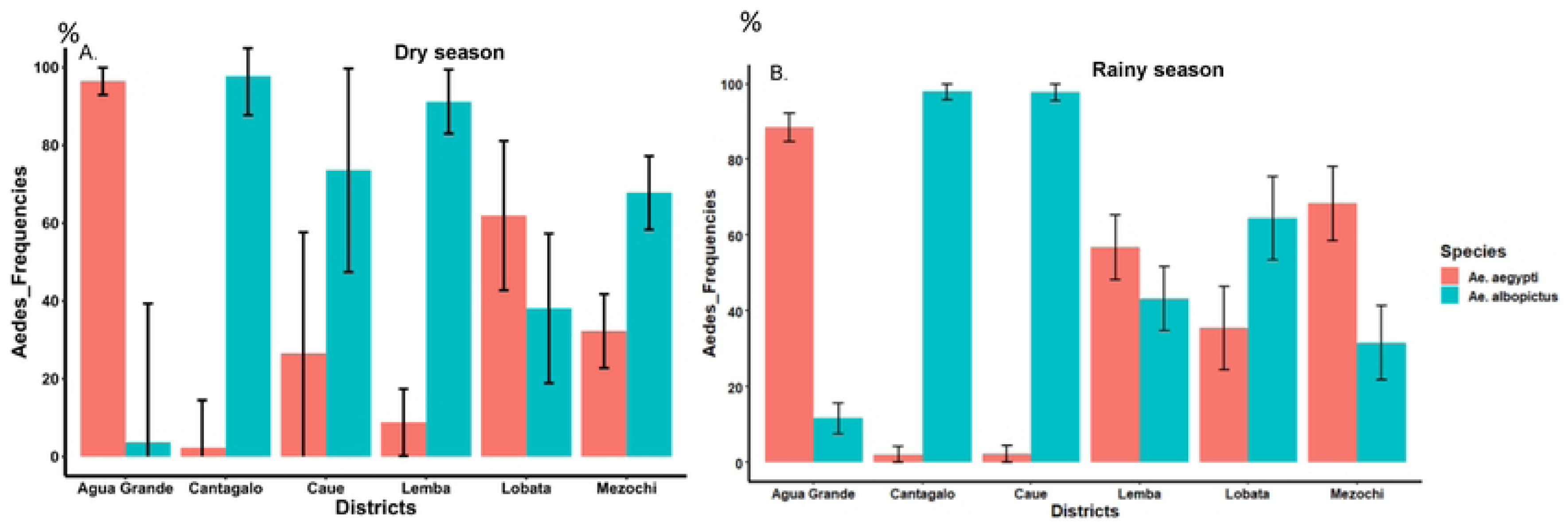
Distribution and prevalence of *Ae. aegypti* and *Ae. albopictus* per health district. A, dry season; B, rainy season.

### Insecticide resistance profile in *Ae. aegypti*

Unfortunately, we successfully obtained only enough *Ae. aegypti* for bioassays. In total 1000 *Ae. aegypti* adults from Sao Tome were tested with seven insecticides. The results revealed that this sample was resistant to DDT (9.2% mortality) and bendiocab (61.4% mortality), and probable resistance to 0.005% alphacypermethrin (97.1% mortality), but was susceptible to 0.25% pirimiphos-methyl, 0.03% deltamethrin, 0.40% permethrin, and fenitrothion (with 100%, 98%, 99% and 100% mortality respectively) (Fig 5).

**Figure 5.**
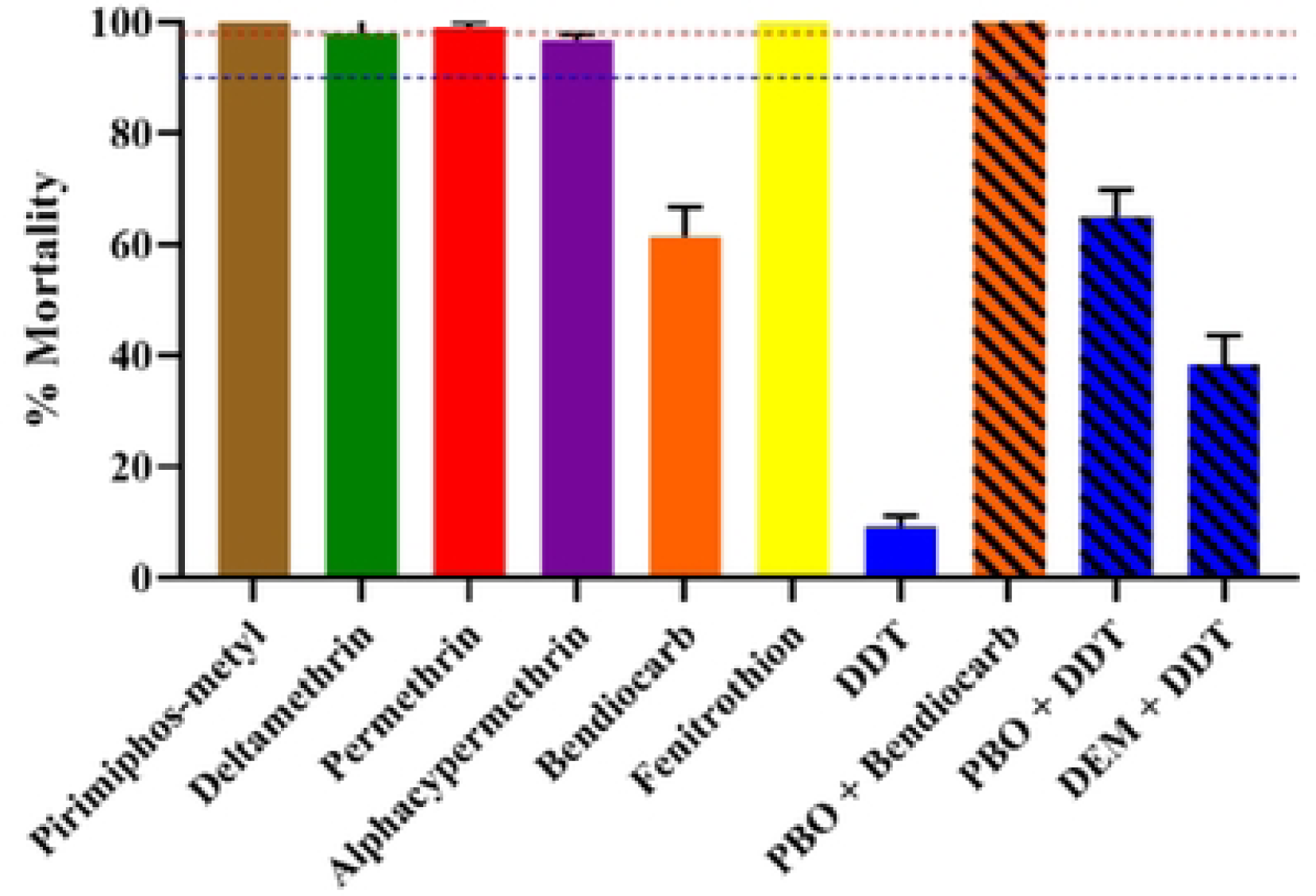
Mortality rates of adult *Aedes aegypti* from Sao Tome when exposed to insecticides alone or with 1 h pre-exposure to synergist. Error bars represent standard error of the mean. PBO Piperonyl butoxide. DEM, diethyl maleate.

### Synergist assay

Results from synergist assays showed a full recovery of susceptibility to bendiocarb after PBO pre-exposure (61.4±5.3 mortality without PBO pre-exposure vs 100.0 ± 0.0% mortality after PBO pre-exposure, *p*<0.0001). However, only a partial recovery to DDT was observed (9.2 ±2.0 mortality without PBO pre-exposure vs 65.0 ±4.7 mortality after PBO pre-exposure, *p*<0.0001) after pre-exposure to PBO and DEM (Fig 3).

### F1534C, V1016I and V410L *kdr* genotyping

Among the 28 specimens of *Ae*. *aegypti* G0 from Sao Tome that were genotyped, no resistant individual was detected with V1016 or V410 mutations, while one heterozygote resistant was found with F1534C mutation.

## Discussion

This was the first study to assess the distribution of *Ae. aegyti* and *Ae. albopictus* in Sao Tome and Principe, the typology of larval habitats, and the susceptibility profile of *Ae. aegypti* to insecticides. The investigations were performed during the first dengue outbreak reported in the country in 2022 [33]. Epidemiological data collected during the outbreak revealed the presence of positive cases of dengue across all the seven health districts of Sao Tome and Principe, however, more than 68% of cases were resident in the health district of Agua Grande [33] which is the most urbanized and most populated area in Sao Tome and Principe.

### Entomological risk and distribution of *Ae. aegypti* and *Ae. albopictus*

The Stegomyia indices calculated in Sao Tome were considerably higher than the thresholds established by WHO for potential dengue and yellow fever transmission [25], in both seasons. Nonetheless, the values of the Breteau index obtained during the rainy season were two-times higher compared to the dry season, indicating that the potential risk for dengue transmission is more pronounced during the rainy season. In general BI was highest in Agua Grande compared to other health districts suggesting the high transmission in this location, which has a higher building density. This observation is supported by the epidemiological data across the country [Á gua Grande (818 cases), Mézôchi (181), Lobata (97), Lemba (20), Caué (23), Cantagalo (47) et RAP (14)] [33]. The entomological data collected during this dengue outbreak show that both *Ae. aegypti* and *Ae. albopictus* are found throughout the country, but geographical and seasonal differences in their distribution were noted. *Ae. albopictus* was found most prevalent in four health districts, whereas *Ae. aegypti* was the predominant species in Agua Grande. This observation is in accordance with the previous data collected in some locations where both species are sympatric. Indeed, studies conducted according to an urbanization gradient have demonstrated that *Ae. aegypti* prefers urban locations with high building density while *Ae. albopictus* is more abundant in rural or peri-urban areas with high vegetation index [34]. This matches with the observation made in different health districts in Sao Tome and Principe.

Furthermore, banana trees were found in almost all habitations in the country. This could explain why, even if *Ae. albopictus* was introduced recently as suggested by [10], this invasive species has become the most predominant species in 4/7 health districts in Sao Tome and Principe. Previous studies in Central Africa have showed that this species tend to replace resident species *Ae. aegypti* in several locations where both are found sympatric [35–37].

### Typology of larval habitats

The entomological investigation revealed that used tyres, discarded tanks and domestic water storage containers were both the most abundant and the most infested larval habitats for *Aedes* spp. in Sao Tome, irrespective of the season; whereas water storage containers were found to be the principal breeding site in Principe. This difference found in the two islands can be explained by deficiencies in the water supply system in Principe, and which is more pronounced compared to Sao Tome, that led residents to store water in containers for a long period. In fact, during the investigations some household owners reported they often store water for a month in drums, bucket and jerrycans. Socio-anthropological studies could help to better elucidate this phenomenon. The high presence of used tyres and discarded tanks positive for immature stages of *Ae. aegypti* and *Ae. albopictus* is in accordance with data previously found in other Central African cities such as Bangui in the Central African Republic [13], Brazzaville in the Republic of the Congo [38], Yaoundé and Douala in Cameroon [12, 14, 20, 36]. On the other hand, the situation in Principe is closer to what has generally found in Southeast Asia where water storage containers are the most prevalent and productive larval habitats for *Ae. aegypti* [39]. In general, these observations suggest that a good waste management system, recycling of used tyres, and communication to support behaviour change could help to reduce breeding sites and the density of *Aedes* in the country. Indeed, during our field surveys discarded containers found were destroyed through perforation, saucers of flower pots were perforated, large water storage containers (cement tanks) were treated using *Bacillus thuringiensis israelensis* (*Bti*) (Fig 2) and advice was given to the population on how to avoid or eliminate *Aedes* larval habitats. This larvicide was chosen because it’s highly specific to Diptera, might be considered a biological control agent and its effectiveness was reported to control dengue vectors [40–42]. In addition, *Aedes* in central Africa have consistently been found susceptible to *Bti* [17, 21, 43]. *Bti* was also readily available on the islands as it was already being used for *Anopheles* larval control by the malaria elimination programme.

### Insecticide susceptibility profile in *Ae. aegypti*

This first study in Sao Tome revealed that *Ae. aegypti* was resistant to bendiocab and DDT, but remains susceptible to permethrin, alphacypermethrin, deltamethrin, pirimipho-methyl and fenitrothion. These findings suggest that all the insecticides tested, except for the two resistant compounds, could be recommended to control *Ae. aegypti* in this country. A decreasing susceptibility of *Ae. aegypti* population from Central Africa towards DDT, notably in Brazzaville and Yaoundé, was already described in 1970s [44], suggesting that this resistance may reflect a continuing selection pressure on *Aedes* populations as suggested previously [43]. Indeed, during the last decade DDT resistance has repeatedly been reported in *Ae. aegypti* [14, 19, 20, 43, 45, 46]. A loss of sensitivity was also observed to bendiocarb with moderate level of resistance. Similar results were recently found in Cameroon [22] and Burkina Faso [47, 48] in Africa, and several countries outside Africa such as Malaysia [46], Colombia [49], and Mexico [50]. The source of selection driving the observed resistance to DDT and bendiocarb in *Ae. aegypti* populations remains unclear, as the programmatic use of insecticides against *Aedes* is limited in the African region [18]. As suggested previously [14, 43], domestic use of insecticides through indoor spraying and impregnated bed nets, and agricultural use could be the main sources of resistance selection in *Aedes* vectors in Central Africa, whilst the use of pesticides in agriculture for the protection of crops could promote the emergence of resistance in mosquitoes through contamination of breeding sites and resting places of mosquitoes [14]., Even though loss of susceptibility in *Aedes* vectors has been reported in other location in central Africa [18–20], *Ae. aegypti* collected from Sao Tome showed a good level of susceptibility to both type I and II pyrethroids tested.

A full recovery of susceptibility to bendiocarb was observed in *Ae. aegypti* from Sao Tome after pre-exposure to PBO synergist suggesting that the cytochrome P450 monooxygenases are playing the main role in the observed resistance. On the other hand, only partial recovery of sensitivity to DDT was seen, both after pre-exposure to PBO or DEM synergists. This observation suggests a possible implication of both cytochrome P450 monooxygenases and glutathione S-transferases in DDT-resistant *Ae. aegypti* as suspected previously [51]. Among the three *kdr* mutations 1534, 410 and 1016 genotyped only one specimen was found to possesses the 1534C allele, indicating that this mutation is not currently involved in pyrethroid/DDT resistance in *Ae. aegypti* in Sao Tome. Nevertheless, these mutations are known to be widely distributed in *Ae. aegypti* [30] and have previously been detected in samples from Central Africa [19, 20]. Due to limitations of sample size, this study was unable to assess insecticide susceptibility in *Ae. albopictus*. Yet it could be useful to extend this study to examine this, since *Ae. albopictus* was found to be most prevalent in four health districts in Sao Tome and Principe, as well as to continue to monitor insecticide resistance in both species across the country.

## Conclusions

This study has provided for the first time the typology of *Aedes* larval habitats and the distribution of *Ae. aegypti* and *Ae. albopictus* across Sao Tome and Principe. The results revealed that these species bred mainly in used tyres, discarded tanks and water storage containers. A pattern of susceptibility to insecticides in *Ae. aegypti* was established enabling this country to quickly implement insecticide-based control interventions in case of future outbreaks. Indeed, organophosphates notably pirimiphos-methyl was used to control *Aedes* adults in Sao Tome and Principe during this dengue outbreak. Findings generated in this study helped to give advice to the population on the practical actions to limit the proliferation of *Aedes* and the Ministry of Health to implement an efficient strategy to control dengue vectors in Sao Tome and Principe. Continued future engagement with political leaders and local communities will also be key to improve systems of water supply and waste management, reducing the number of potential breeding sites in the domestic and peri-domestic environment to reduce the risk of future outbreaks. As there is currently a lack of control programs against arbovirus in many countries in Africa, we recommend to use an integrated vector control strategic approach to benefit from the successes of well-established vector control program against malaria, enabling more efficiencies in the fight against mosquito-borne diseases. This approach will permit to combat mosquitoes in general to reduce the risk of mosquito-borne diseases transmission.

## Acknowledgements

We would like to thank the populations from the different collection sites for their cooperation during mosquito sampling.

## Authors’ contributions

BK, JA, VS, and AA conceived and designed the experiments. BK and JA participated in mosquito collections. BK, CK and AY performed the bioassays. BK, TA and SC carried out the data analyses. CK and AY conducted the molecular analyses. BK wrote the paper. All authors read and approved final version of the manuscript.

## Notes

### Competing Interest Statement

The authors have declared no competing interest.

